# Characterization of a late blight resistance gene homologous to *R2* in potato variety Payette Russet

**DOI:** 10.1101/2020.09.27.315812

**Authors:** Hari S. Karki, Dennis A. Halterman, Jeffrey B. Endelman

## Abstract

Breeding for late blight resistance has traditionally relied on phenotypic selection, but as the number of characterized resistance (R) genes has grown, so have the possibilities for genotypic selection. One challenge for breeding russet varieties is the lack of information about the genetic basis of resistance in this germplasm group. Based on observations of strong resistance by ‘Payette Russet’ to genotype US-23 of the late blight pathogen *Phytophthora infestans* in inoculated experiments, we deduced the variety must contain at least one major R gene. To identify the gene(s), 79 F1 progeny were screened using a detached leaf assay and classified as resistant vs. susceptible. Linkage mapping using markers from the potato SNP array revealed a single resistant haplotype on the short arm of chromosome group 4, which coincides with the *R2/Rpi-abpt/Rpi-blb3* locus. PCR amplification and sequencing of the gene in Payette revealed it is homologous to *R2*, and transient expression experiments in *Nicotiana benthamiana* confirmed its recognition of the *Avr2* effector. Sequencing of a small diversity panel revealed a SNP unique to resistant haplotypes at the *R2* locus, which was converted to a KASP marker that showed perfect prediction accuracy in the F1 population and diversity panel. Although many genotypes of *P. infestans* are virulent against *R2*, even when defeated this gene may be valuable as one component of a multi-genic approach to quantitative resistance.

## INTRODUCTION

One of the most significant pathogens of potato (*S. tuberosum* L.) is the oomycete *Phytophthora infestans*, which causes late blight. In the USA and EU, growers typically apply fungicides for late blight control upwards of 10 times per season, at a cost of US$500 per hectare (Johnson et al. 2000; Guenthner et al. 2001; Haverkort et al. 2008). Late blight also affects tubers and can render the harvested crop unmarketable from the field or following storage when susceptible varieties are left unprotected. To mitigate the economic impact of late blight, breeders have spent decades introgressing disease resistance into cultivated potato from wild relatives. Because of the autotetraploid (2n = 4x = 48) genome of cultivated potato, however, this resistance is generally not fixed and must be selected in each generation. The selection process is more efficient, especially considering that many thousands of new clones are created by a single breeding program each year, when using genetic markers instead of via inoculated disease assays. Genetic markers are also critical when stacking multiple resistance genes (targeting different effectors in the pathogen) in a single variety.

A number of North American varieties have been developed with some degree of late blight resistance, but it is only recently that significant progress has been made toward identifying the genetic basis. One breakthrough was the potato SNP array (Hamilton et al. 2011; Felcher et al. 2012) combined with new software for genetic mapping (Hackett et al. 2014), which Massa et al. (2015) used to map the R gene in ‘Jacqueline Lee’ to the long arm of chromosome 9. This coincided with the published location of *R8* (Jo et al. 2011), leading to the hypothesis that *R8* was responsible for the resistance, which was confirmed by Vossen et al. (2016). Another breakthrough was the development of DNA bait libraries based on all predicted NB-LRR genes in the potato genome (Jupe et al. 2013). This method, called RenSeq, relies on hybridization of genomic DNA with the bait library to isolate candidate genes for more economical sequencing. Although originally used for mapping unknown R genes, it can also be used to detect previously characterized R genes (Armstrong et al. 2019).

The first aim of the current project was to identify the genetic basis of resistance in ‘Payette Russet’, which was released as a fry processing variety in 2015 and publicized as late blight-resistant based on inoculated field trials in Oregon and Idaho (Novy et al. 2017). Our interest was piqued after observing almost no injury to Payette in a whole-plant assay with *P. infestans* genotype US-23, while the other potato genotypes in the experiment were largely destroyed. In addition to isolating and characterizing the R gene in Payette, we developed a KASP marker that can be used for breeding.

## MATERIALS AND METHODS

### Plant material and late blight assay

The F1 population W16215 was generated by crossing Payette Russet with A0012-5 (GemStar Russet × Wallowa Russet). A Wisconsin isolate of *P. infestans* genotype US-23 was used to screen both parents and the F1 population (grown from seed tubers in a greenhouse). The isolate was maintained on Rye A media at 18°C and refreshed periodically. Sporangia were harvested from 12–14 day-old culture plates by flooding with ice cold water and then kept at 4°C for 2–4 hours to release zoospores. The detached leaf assay protocol of Vleeshouwers et al. (1999) was followed with some modifications. Briefly, compound leaves having at least three leaflets were collected from 5–8 week-old plants into Nunc assay plates with a moist paper towel. The abaxial side of each leaflet was inoculated (4 to 6 droplets per leaflet) with 10-μl droplets of zoospores (50,000 zoospores per ml), and samples were stored in an incubator at 16°C and 16 h daylight. Disease was scored 7 days after inoculation using a 1–9 scale (Karki et al. 2020), and offspring were ultimately classified as resistant (scores 7–9) vs. susceptible (scores 1–4) after three replications.

### SNP array genotyping and genetic mapping

Both parents and the F1 population were genotyped at the University of Minnesota using version 3 of the potato SNP array. Genomic DNA was extracted with the DNeasy Plant Maxi Kit (QIAGEN, Hilden, Germany) according to the manufacturer instructions. Genotype calls were made using the fitPoly R package (Voorrips et al. 2011; Zych et al. 2019). R package MAPpoly (Mollinari et al. 2020) was used to phase the parental genotypes and calculate identity-by-descent (IBD) probabilities for each offspring based on the SNP positions in version 4.03 of the DM reference genome (Potato Genome Sequencing Consortium 2011; Sharma et al. 2013). R packages diaQTL (available at https://github.com/jendelman/diaQTL) and BGLR (Pérez and de los Campos 2014) were used to map the binary resistance phenotype (R/S) via Bayesian probit regression on the IBD probabilities. The LOD score for each marker was calculated as the difference in log-likelihood (evaluated at the posterior mean) between the QTL and no-QTL model. The predictive ability of each marker was quantified based on the phi-correlation coefficient for the 2×2 contingency table between the predicted and observed responses.

### R gene isolation and validation in *N. benthamiana*

The primers used to isolate *Rpi-blb3* (Lokossou et al. 2009) were modified (Table S1) to amplify the R gene in Payette by including additional 5′ and 3′ overhangs to facilitate cloning into custom USER expression vectors (kindly provided by The Sainsbury Laboratory, Norwich, UK). The gene was amplified by PCR in a 25 μl reaction with initial denaturation of 95°C for 3 min, 30 cycles of [98°C for 20 sec, 65°C for 30 sec, and 72°C for 3 min], and final extension of 72°C for 10 min using KAPA master mix (Roche). PCR products were visualized using a 0.8% agarose gel, then excised and purified using the Zymoclean DNA recovery kit (Zymo Research) according to the manufacturer’s instructions. The purified DNA fragment (40 ng) was ligated into pICSLUS0003OD (35S USER vector) with the USER enzyme (New England Biolabs). After verifying the presence of the insert by colony PCR, purified plasmids from two clones were submitted for next-generation sequencing and contig assembly (using SPAdes) at the Purdue Genomics Core Facility (Purdue University, West Lafayette, IN). Contigs were mapped against the reference sequence *Rpi-abpt* (GenBank Accession No. FJ536324.1) using Geneious Prime® 2019.0.4.

To verify the functionality of the R gene isolated from Payette Russet, purified plasmid was used for transformation of *A. tumefaciens* strain GV3101. In the first experiment, overnight culture of *Agrobacterium* was centrifuged, washed, and suspended in MMA media (Abdullah and Halterman 2018) to OD_600_ = 0.2, followed by infiltration of *N. benthamiana* leaves (collected after 4–5 weeks of growth at 26°C and 12 h daylight) with a needleless 1 mL syringe. *Agrobacterium* culture containing *Rpi-abpt* (kindly provided by Jack Vossen, Wageningen University & Research) was used as a positive control. 24 h after infiltration, a detached leaf assay using *P. infestans* genotype US-23 was performed as described above. For the second experiment, equal volumes of *Agrobacterium* culture containing the R gene and effector *Avr2* (Gilroy et al. 2011; kindly provided by Jack Vossen, Wageningen University & Research) were mixed after centrifugation, washing, and suspension in MMA media to OD_600_ = 0.4. *N. benthamiana* leaves were infiltrated with the mixed culture and evaluated after 6 days. The cell death-inducing elicitin INF1 was used as a positive control (Kamoun et al. 1998).

### KASP marker design

To identify a SNP unique to resistant haplotypes at the *R2/Rpi-abpt/Rpi-blb3* locus, the primers (Table S1) and PCR amplification protocol described above were applied to a larger set of varieties. PCR products (Fig. S1) were purified using the Zymoclean DNA recovery kit (Zymo Research) and submitted (without cloning into plasmids) for next-generation sequencing and contig assembly at the Purdue Genomics Core Facility (Purdue University, West Lafayette, IN). Contigs were mapped against the *Rpi-abpt*^T87^ haplotype from Payette Russet and aligned using Geneious Prime® 2019.0.4. 50 bp of flanking sequence on either side of the target SNP was submitted to LGC Genomics (MA, USA) for KASP marker design and validation.

## RESULTS

### Payette contains a homolog of *R2*

An F1 population containing 79 progeny of Payette Russet was screened for resistance to the US-23 genotype of *P. infestans* via detached leaf assay. Clear visual differences (Fig. 1) were observed for resistant (R) vs. susceptible (S) phenotypes to allow for binary classification. The ratio 36R:43S was not significantly different than 1:1 (*χ*^2^ = 0.62, *p* = 0.4), suggesting one R gene with simplex dosage. The population was genotyped with version 3 of the potato SNP array, which generated 11,094 polymorphic markers. Genetic mapping confirmed the presence of a single resistant haplotype on the short arm of chromosome group 4 (Fig. 2).

**Figure 1.**
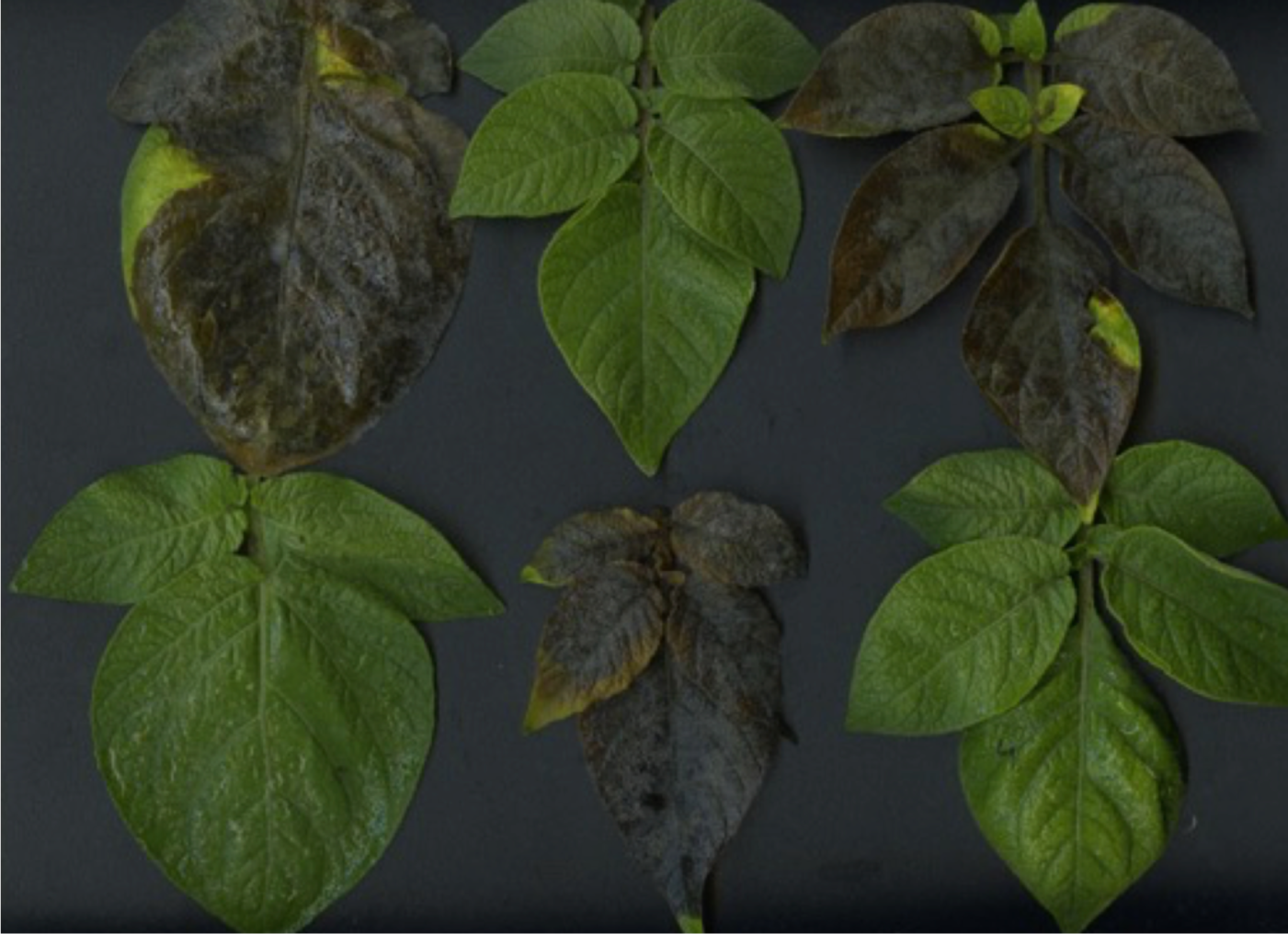
Detached leaf assay for the W16215 (A0012-5 x Payette Russet) mapping population. Photograph illustrates the appearance of resistant vs. susceptible progeny 7 days after infection with a Wisconsin isolate of the *P. infestans* genotype US-23.

**Figure 2.**
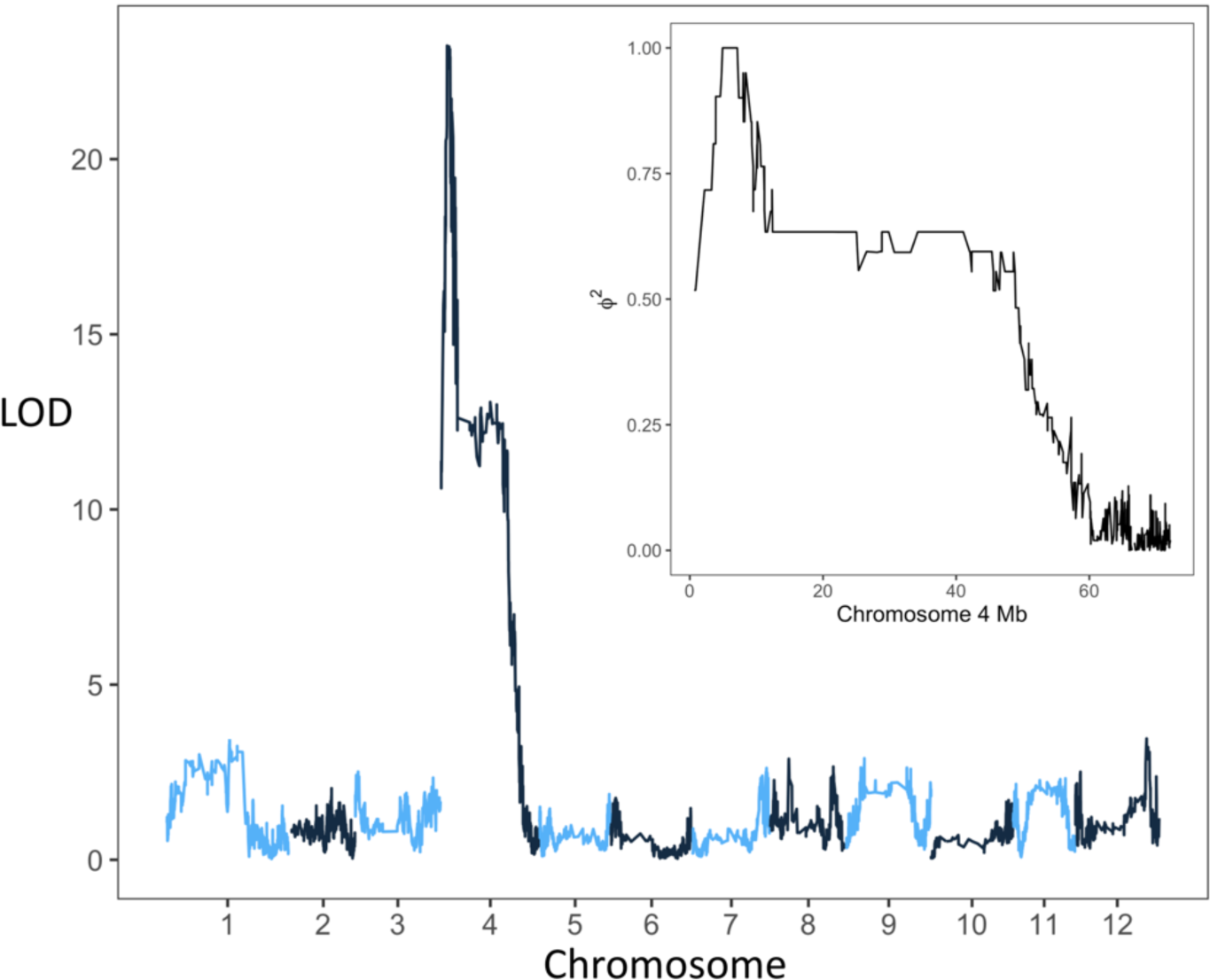
LOD profile based on Bayesian probit regression of the binary resistance trait against parental haplotype probabilities. The peak on chromosome 4 is due to a single haplotype in Payette Russet. Inset: Phi-squared correlation showing the perfect correlation (= 1) on the short arm of chromosome 4.

Based on this location, we hypothesized the R gene to be homologous to *R2, Rpi-abpt*, and *Rpi-blb3* (Li et al. 1998; Park et al. 2005a; Park et al. 2005b; Lokossou et al. 2009). The primers for *Rpi-blb3* generated an amplicon of the expected size (∼2.5 kb), which was cloned into a plasmid for sequencing. Two clones were sequenced, and both were identical to the published sequence for *Rpi-abpt* except for a synonymous C→T substitution at position 87 (relative to the start of the coding sequence). This matches the haplotype in Pentland Dell discovered by Armstrong et al. (2019), who reported the C→T substitution at position 86 instead of 87 based on the bp numbering convention of the BED file format (I. Hein, personal communication).

*N. benthamiana* was used to verify the functionality of the *R2* homolog isolated from Payette. Leaves infiltrated with *Agrobacterium* harboring the gene showed no signs of late blight when inoculated with *P. infestans* genotype US-23 (Fig. 3A & B), and co-infiltration with *Agrobacterium* culture harboring *Avr2* resulted in a hypersensitive response (Fig. 3C), thereby confirming recognition of this effector.

**Figure 3.**
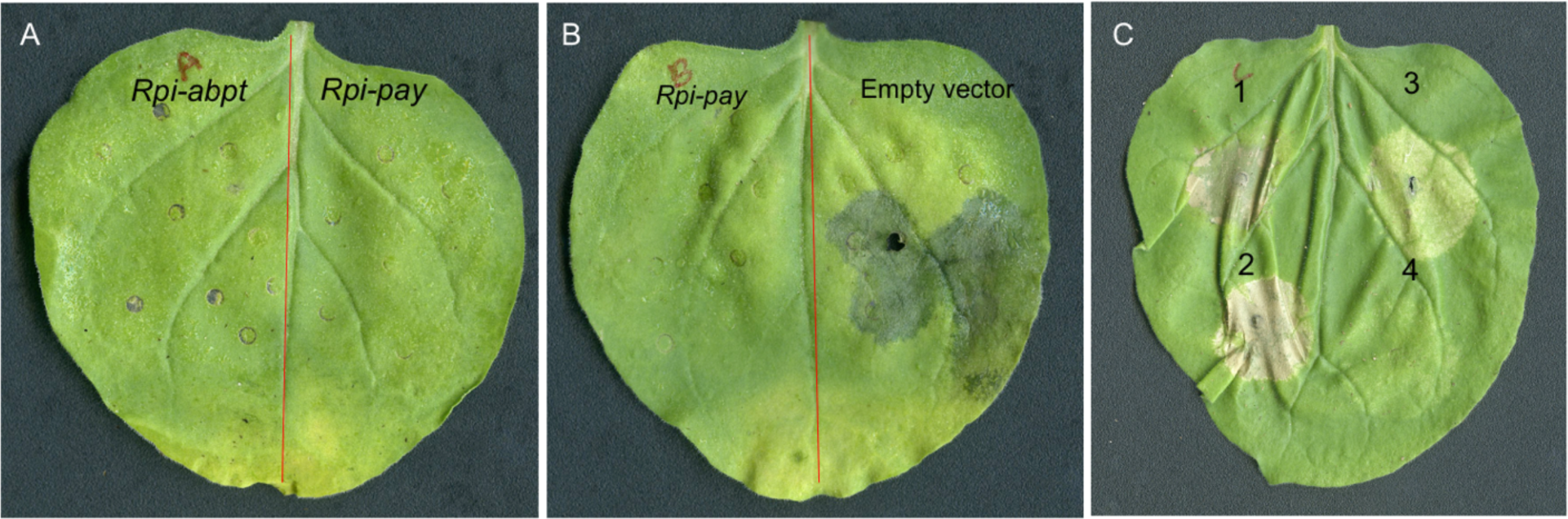
Genetic complementation and effector recognition assay in *N. benthamiana* leaves. (A & B) Transient expression of *Rpi* genes followed by inoculation with *P. infestans* genotype US-23. *Rpi-abpt* and empty vector were used as a positive and negative control, respectively. Photographs were taken 6 days after inoculation. (C) *Agrobacterium* mediated co-expression of the *Avr2* effector and *Rpi* genes. Infiltration points correspond to (1) the cell death-inducing elicitin *INF1*; (2) Payette R gene + *Avr2*; (3) *Rpi-abpt* + *Avr2*; and (4) Payette R gene alone. Photographs were taken 6 days after co-infiltration.

### Design and application of a KASP marker

To identify a SNP unique to resistant haplotypes at the *R2/Rpi-abpt/Rpi-blb3* locus, the same primers used to isolate the *Rpi-abpt*^T87^ haplotype from Payette were applied to a diverse set of potato varieties (Saginaw Chipper, Jacqueline Lee, Palisade Russet, ND028673B-2Russ, A0012-5, Red Prairie, and Rio Colorado) that, based on their susceptible phenotype in the detached leaf assay, were presumed to lack a functional *R2* homolog. PCR products were purified and submitted for next-generation sequencing. The number of gene-length contigs that mapped to the *Rpi-abpt*^T87^ sequence ranged from 0 to 5 per variety (Fig. 4). From Payette Russet, which was used as a positive control in the experiment, a second haplotype was discovered with a 420 bp insertion and premature stop codon. This second haplotype was not discovered during the initial sequencing, presumably because of the small number of clones that were sequenced. Based on visual inspection of the multiple sequence alignment, we noticed that the resistant haplotypes (first five in Fig. 4) all contained C at 2223 bp (from the beginning of the coding sequence for *Rpi-abpt*), while the susceptible haplotypes contained T, making it an ideal SNP for breeding.

**Figure 4.**
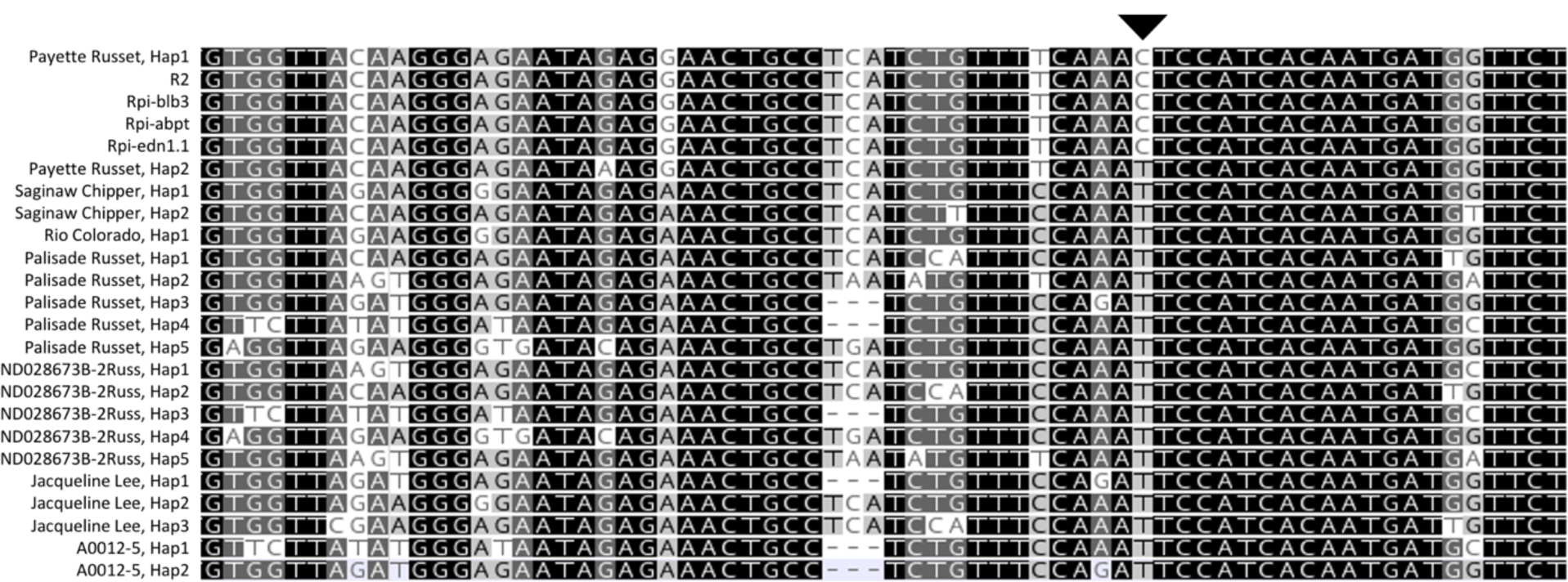
Resistant (rows 1–5) and susceptible (rows 6–24) haplotypes aligned to the *Rpi-abpt*^T87^ haplotype discovered in Payette Russet (“Hap1”). Inverted triangle indicates the C/T SNP used for KASP marker development.

The C/T SNP and flanking sequence were submitted for KASP marker design (Table 1) and genotyping of the small diversity panel and F1 population used for mapping. Samples were also genotyped using the *R2* KASP marker published by Meade et al. (2020). Both KASP markers had a perfect correlation with the resistance phenotype in the F1 population and correctly predicted the absence of an *R2* homolog in the diversity panel.

**Table 1.**
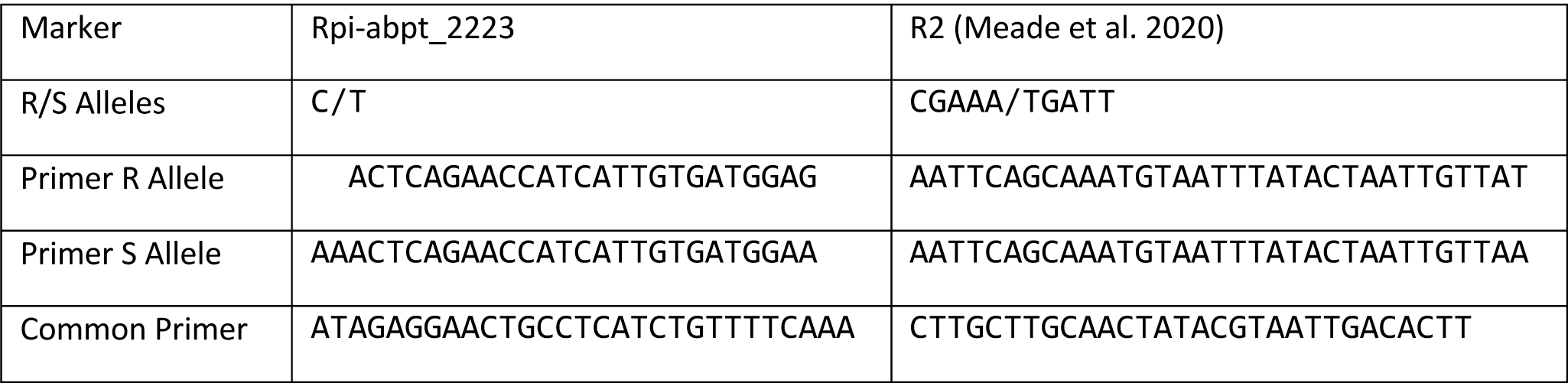
KASP markers for the *R2/Rpi-abpt/Rpi-blb3* locus.

## DISCUSSION

Our results demonstrate that *Rpi-abpt*^T87^ provides strong resistance to the US-23 genotype of *P. infestans*, which is currently the most common genotype in the USA, and this resistance is available in an elite russet background. How will this impact breeding? Although there is a long history of using *R2* homologs in Europe, it is no longer a primary target because many isolates of *P. infestans* are virulent against it. As mentioned already, the functional Payette haplotype is identical to that in Pentland Dell, which was released in Great Britain in 1963 and promoted as having durable resistance due to the presence of three R genes (*R1, R2, R3*). This contributed to its rapid adoption, becoming the third most popular variety by 1968, but widespread late blight on Pentland Dell that year shattered the illusion of durable resistance (Malcolmson 1969). Pilet et al. (2005) reported that, prior to the large scale deployment of cultivars containing *R2* homologs in western France, the onset of late blight on those cultivars was significantly later compared to susceptible varieties, and most *P. infestans* isolates were avirulent. However, these advantages sharply decreased as the acreage of *R2*-containing varieties increased. Even within the USA, the avirulence of US-23 against *Rpi-abpt*^T87^ may be somewhat anomalous, as most of the formerly dominant *P. infestans* genotypes, such as US-8, were reported to be virulent against *R2* (Goodwin et al. 1995). Due to the scarcity of sexually compatible mating types in the US, however, *P. infestans* populations tend to evolve more slowly and may have difficulty overcoming *Rpi-abpt*^T87^ if it becomes more widely deployed.

One response to the perennial tug-of-war between *P. infestans* adaptation and resistance breeding has been to avoid major R genes and select for quantitative late blight resistance in the field (Wastie 1991). This is challenging in the context of current best practice for breeding, which emphasizes rapid parent selection driven by genomic prediction (Slater et al. 2014; Cobb et al. 2019), because hundreds of clones would need to be evaluated each year to maintain accuracy of the prediction model. Such trials are prohibited in some states (such as Wisconsin), and even when they are allowed, require significant resources. Our view is that stacking R genes is still valuable, and readily accomplished with genomic selection, since even defeated R genes can contribute to quantitative resistance (Stewart et al. 2003; St. Clair 2010). For many programs, the goal of breeding for late blight resistance is to delay when preventive fungicide applications begin or reduce their frequency, not completely eliminate them. Viewed in this conext, our discovery of an *R2* homolog in Payette can benefit russet variety development.

## Acknowledgments

Funding for this research was provided by the Wisconsin Potato and Vegetable Growers Association, the USDA NIFA/NSF Plant Biotic Interactions Program Award 2018-67014-28488, and PepsiCo. We thank Grace Christensen and Peyton Sorensen for their assistance in the lab and greenhouse, as well as personnel at the UW Rhinelander Agricultural Research Station.

## Supplemental Information

**Table S1.**
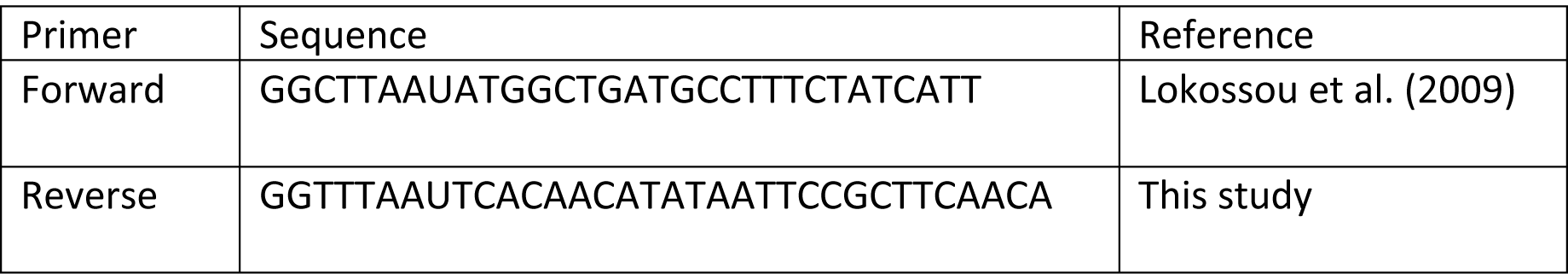
Primer sequences used to amplify *R2* homologs.

**Fig S1.**
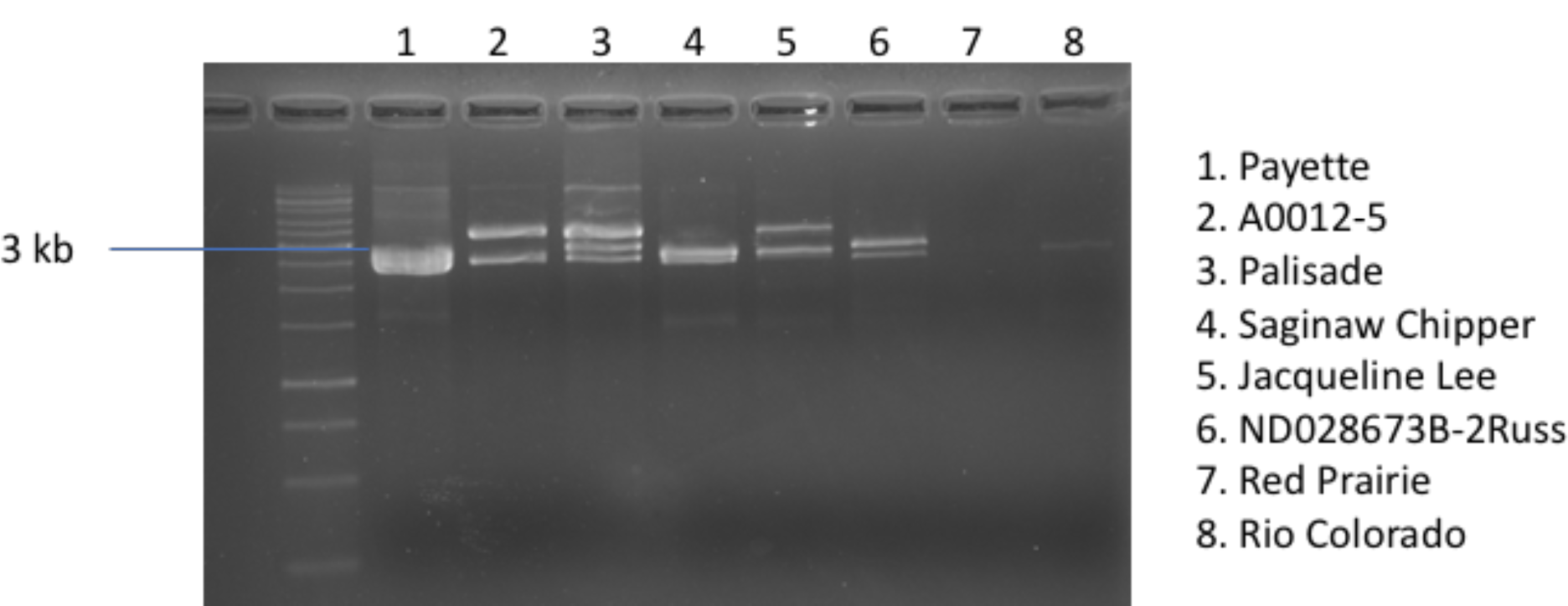
Gel electrophoresis of PCR products using the primers in Table S1.

